# Correlation Analysis of Breast Cancer Molecular Characteristics and Epithelial-Mesenchymal Transition in Circulating Tumor Cells: Based on Clinical Cases

**DOI:** 10.1101/2025.07.08.663630

**Authors:** Jiayu Guan, Qianqi Chenli, Xiaoyu Pu, Wenbin Zhou

**Affiliations:** The Second Clinical Medical College, Jinan University, Guangdong, Shenzhen 518020, China; Department of Breast Surgery, Shenzhen People’s Hospital, Guangdong, Shenzhen 518020, China; SurExamBio-TechCo.,Ltd

**Keywords:** Breast cancer, molecular subtype, circulating tumor cells, epithelial-mesenchymal transition

## Abstract

**Background:** This study aimed to investigate the correlation between epithelial-mesenchymal transition (EMT) in circulating tumor cells (CTCs) from peripheral blood and molecular characteristics of breast cancer.

**Methods:** We enrolled 48 breast cancer patients who underwent CTC testing at Shenzhen People’s Hospital’s Breast Surgery Department between September 2022 and May 2024. Intraoperative tissue samples underwent routine pathological and immunohistochemical examination. CTCs were detected using Canpatrol™ technology. Clinical data were analyzed statistically using SPSS 27.0. Categorical data were presented as percentages/frequencies. Comparisons used chi-square tests, with Yates correction for expected frequencies <5 but ≥1. Statistical significance was set at P<0.05 (α=0.05).

**Results:** EMT status showed no significant differences between ER-positive/negative, PR-positive/negative, HER2-positive/negative, Ki-67 ≥14%/<14%, tumor size, or lymph node metastasis groups (P>0.05). Luminal-type tumors exhibited lower EMT grades than other subtypes, while triple-negative breast cancer (TNBC) showed higher EMT grades (P<0.05).

**Conclusion:** EMT grade correlates with luminal-type and TNBC subtypes, suggesting predictive value for these breast cancer classifications.

## Introduction

Breast cancer is one of the most common malignant tumors among women worldwide (1, 2), characterized by high heterogeneity, with recurrence and metastasis being the primary causes of patient mortality (3). The molecular subtypes of breast cancer—Luminal, HER2-overexpressing, and triple-negative—significantly influence prognosis and treatment strategies. The highly heterogeneous molecular characteristics and complex biological behaviors of breast cancer have long presented significant challenges in clinical diagnosis and treatment. In recent years, circulating tumor cells (CTCs), considered the “seeds” of distant metastasis, have shown that their epithelial-mesenchymal transition (EMT) process is closely associated with tumor invasion and metastatic potential (4-6). Studies (7) indicate that CTCs from different molecular subtypes of breast cancer (such as Luminal, HER2-positive, and triple-negative) exhibit distinct EMT phenotypic characteristics, offering new insights into breast cancer metastasis mechanisms (8).

Recent studies show that CTCs in triple-negative breast cancer display a pronounced mesenchymal phenotypic shift, whereas the Luminal subtype tends to maintain more epithelial traits (9, 10). This study systematically analyzed the EMT profiles of CTCs in the peripheral blood of patients with different molecular subtypes of breast cancer, revealing a correlation between EMT progression stages and both the Luminal subtype and TNBC. These findings reveal the intrinsic link between molecular heterogeneity and metastatic potential in breast cancer, establishing a critical foundation for further investigation of the clinical translation potential of EMT markers.

## Methods

Study Subjects: We enrolled 48 histopathologically confirmed breast cancer patients treated at the Department of Breast Surgery, Shenzhen People’s Hospital (September 2022–May 2024) who underwent CTC testing. Postoperative breast masses underwent routine paraffin sectioning and immunohistochemical analysis in the pathology department. Two pathologists performed pathological and immunohistochemical evaluations. Postoperative data including pathological type, tumor size, lymph node metastasis count, ER, PR, HER2, and Ki-67 status were collected for statistical analysis.

Statistical Analysis: Clinical data were analyzed using SPSS 27.0. Categorical data were expressed as percentages and frequencies. Comparisons used χ^2^ tests, with continuity-corrected χ^2^ applied for theoretical frequencies <5 and ≥1. Statistical significance was set at P<0.05 (α=0.05).

Grouping: ER-positive (>1% estrogen receptor-positive tumor cells); PR-positive (>1% progesterone receptor-positive cells). HER2 (3+) denoted positivity; HER2 (1+) denoted negativity. HER2 (2+) cases required FISH confirmation. Ki-67 ≥14% indicated high expression; <14% indicated low expression. Axillary lymph node metastasis (≥1 node) was considered positive. Tumors were stratified by 2 cm size cutoff. EMT ratio = (mixed + mesenchymal types)/total cases. EMT status was defined as positive (>75% ratio) or negative (≤75%).

CTC Detection: The CanPatrol™ CTC assay (Surexam) involved:

1. Isolation: 10 mL peripheral blood underwent RBC lysis, followed by size-based nanotechnology enrichment of CTCs.
2. Typing/Identification: A novel multiple-mRNA in situ analysis (MRIA) localized specific genes in CTCs. Simultaneous RNA probe hybridization with target genes, coupled with fluorescence signal amplification, achieved single-copy mRNA detection sensitivity.

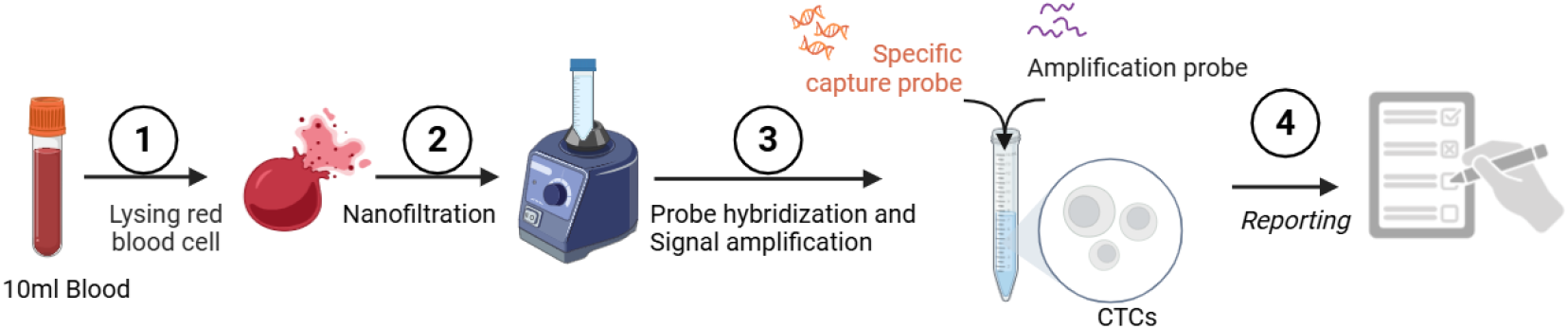

## Results

1. Relationship between EMT grade and pathological characteristics As shown in Table 1, there were no statistically significant differences in EMT grade among ER/PR-positive and ER/PR-negative patients, HER-2-positive status, Ki-67 levels ≥14% vs. <14%, tumor size, or lymph node involvement (P > 0.05).
2. Relationship between EMT grade and molecular subtypes of breast cancer

**Table 1.** Comparison of pathological features with positive and negative EMT (%)

| Molecular typing | Grouping | Number of cases n=48 (%) | EMT positive n=16 (%) | EMT negative n=34 (%) | $\chi^2$ | P |
| --- | --- | --- | --- | --- | --- | --- |
| ER | Positive | 34 | 11 | 23 | 0.000 | 1.000 |
|  | Negative | 14 | 5 | 9 |  |  |
| PR | Positive | 31 | 10 | 21 | 0.046 | 0.831 |
|  | Negative | 17 | 6 | 11 |  |  |
| Her-2 | Positive | 10 | 1 | 9 | 1.911 | 0.167 |
|  | Negative | 38 | 15 | 23 |  |  |
| Ki-67 | $< 14\%$ | 15 | 5 | 10 | 0.000 | 1.000 |
| | $\geq 14\%$ | 33 | 11 | 22 | | |
| Axillary lymph nodes | Positive | 14 | 6 | 8 | 0.315 | 0.575 |
|  | Negative | 34 | 10 | 24 |  |  |
| Tumor size | $\leq 2\text{cm}$ | 35 | 12 | 23 | 0.000 | 1.000 |
| | $> 2\text{cm}$ | 13 | 4 | 9 | | |

As shown in Table 2, luminal-type tumors had lower EMT grades than other subtypes, whereas triple-negative breast cancer (TNBC) showed higher EMT grades, with statistical significance (P< 0.05).

**Table 2.** Comparison of positive and negative EMT (%) in Luminal, HER-2 overexpressing, and TNBC patients.

| Molecular typing | Grouping | Number of cases n=48 (%) | EMT positive n=21 (%) | EMT negative n=27 (%) | $\chi^2$ | P |
| --- | --- | --- | --- | --- | --- | --- |
| Luminal type | Yes | 34 | 11 | 23 | 6.153 | 0.013 |
|  | No | 14 | 10 | 4 |  |  |
| HER-2 overexpression type | Yes | 5 | 1 | 4 | 0.429 | 0.513 |
|  | No | 43 | 20 | 23 |  |  |
| TNBC | Yes | 9 | 9 | 0 | 11.567 | $< 0.001$ |
|  | No | 39 | 12 | 27 |  |  |

## Discussion

The metastatic process of breast cancer involves the detachment of circulating tumor cells (CTCs) from the primary tumor and their entry into the bloodstream (11), with epithelial-mesenchymal transition (EMT) being one of the key mechanisms (12). EMT allows epithelial cells to adopt a mesenchymal phenotype, enhancing their migratory and invasive capabilities while promoting stemness characteristics (13, 14). Research has shown that CTCs exhibit epithelial, mesenchymal, and hybrid phenotypes, with mesenchymal CTCs associated with poorer prognosis (15, 16). Differences in EMT characteristics exist among molecular subtypes of breast cancer: triple-negative breast cancer (TNBC) is more likely to display a mesenchymal phenotype and worse prognosis (17). In such cases, combining traditional chemotherapy with drugs targeting mesenchymal CTCs may help suppress their migration and invasion. Conversely, the hybrid phenotype in HER2-positive subtypes may correlate with resistance to targeted therapy (18, 19). If hybrid CTCs are detected, suggesting resistance to targeted therapy, treatment strategies could be adjusted, such as combining EMT pathway inhibitors with HER2-targeted drugs to enhance efficacy.

This study has several limitations. First, the relatively small sample size and single-center origin may affect the results’ representativeness and generalizability. Future multicenter registry studies are needed to expand sample diversity. Second, although multiple CTC detection technologies exist (20, 21), they vary in strengths and weaknesses, lacking standardized protocols. Currently, there is no international consensus on EMT marker selection, and methodological differences across studies may impact comparability. This study employed the Canpatrol™ CTC detection technology to identify epithelial, mesenchymal, and hybrid CTCs. Additionally, CTCs were assessed at a static time point, which prevented us from dynamically tracking EMT progression and its temporal relationship with treatment response.

Further research into CTC EMT status and the molecular biology of breast cancer holds promise for breakthroughs in diagnosis, treatment, and prognosis assessment (22). Through interdisciplinary collaboration and ongoing clinical exploration, more precise and effective personalized treatment strategies can be developed for breast cancer patients, improving overall treatment outcomes (23).

## Conclusion

In this study, we examined the relationship between EMT grade and tumor size, immunohistochemical (IHC) results, molecular subtypes, and axillary lymph node metastasis status in breast cancer. Our findings demonstrate a correlation between EMT grade and molecular subtypes, with luminal subtypes exhibiting lower EMT grades and TNBC showing higher EMT grades. EMT grade may have predictive value for luminal and TNBC patients. Luminal breast cancer is often associated with better prognosis, while TNBC typically indicates poorer outcomes. Thus, CTC EMT grading may help guide breast cancer treatment and prognostic evaluation.

